# Outside your shell: how temperature shapes genetic variation in two species of congeneric marine snails

**DOI:** 10.1101/2024.09.12.612762

**Authors:** DM Wuitchik, JE Fifer, AK Huzar, JA Pechenik, LH Uricchio, SW Davies

## Abstract

Intertidal organisms withstand extreme temperature fluctuations, and their ability to cope with this variation may affect their distributions across the seascape. Genetic variation and local environments likely interact to determine variation in thermal performance across intertidal species’ ranges, so characterizing the relationship between temperature variation and population structure is key to understanding the biology of marine invertebrates. Here, we use 2bRAD-sequencing to examine population genetic structure in two congeneric intertidal marine gastropods (*Crepidula fornicata*, *C. plana*), sampled from locations along a natural temperature gradient on the Northeast shores of the United States. These two species share similar life histories, yet *C. plana* exhibits a narrower distribution than *C. fornicata*. Our results demonstrate that both species show patterns of genetic divergence consistent with isolation by distance, though this pattern was only significant in *C. fornicata*. Both putatively selected and neutral loci displayed significant spatial structuring in *C. fornicata*; however, only putatively selected loci showed significant clustering in *C. plana*. When exploring whether temperature differences explained genetic differentiation, we found that 9-12% of genetic differentiation was explained by temperature variation in each species even when controlling for latitude and neutral population structure. Our results suggest that temperature shapes adaptive variation across the seascape in both *Crepidula* species and encourages further research to differentiate our results from models of neutral evolutionary drift.

## Introduction

Species distributions are governed by a variety of biotic and abiotic factors. In marine taxa, many species disperse through a bipartite lifecycle, which includes a sessile adult stage and a planktonic larval stage. Historically, it had been assumed that these highly dispersive larvae lead to panmixia between populations, even across large spatial scales (Keller et al., 2022; Levin, 2006). But advancements in modern genomic sequencing technologies for non-model systems have challenged this view and research has demonstrated that many marine organisms are much more genetically structured than previously appreciated (for review, see: Selkoe et al., 2016). This genetic structure can manifest through neutral or selective processes. Neutral processes can structure genetic variation through differences in migration rates (Von Der Heyden et al., 2008) as well as through oceanographic properties that facilitate or constrain larval dispersal across the seascape (Pappalardo et al., 2015). However, even in the face of high gene flow, selection can also shape allele frequencies following environmental clines (Slatkin, 1973). Disentangling how these evolutionary processes operate across environmental gradients is key baseline knowledge for understanding how environments shape species distributions and how these distributions might shift under climate change.

Temperature is an important abiotic factor shaping species distributions as temperatures constrain many physiological processes (Kültz, 2005). Due to its ubiquitous effects, temperature leaves broad fingerprints on the genetic architecture of species and populations (Clarke, 2003). Temperature can act as a selective agent due to its ability to govern the basic functions of numerous cellular processes (Somero, 1995), which can ultimately shape patterns of local adaptation (Sternberg & Thomas, 2014). Perhaps less intuitively, temperature regimes can also drive population genetic patterns via neutral processes. For example, major heat waves can cause large declines in effective population sizes of exposed populations, leading to drift becoming a stronger evolutionary force relative to selection (Lande, 1976). As climate change continues to warm habitats globally (IPCC, 2022), these warmer temperatures will be a critical factor determining genomic diversity and structure both within and across species in the coming decades.

Temperatures naturally vary along latitudinal gradients and this variation can be leveraged to investigate how populations will respond to temperature changes (De Frenne et al., 2013). For example, latitudinal gradients can be used to generate and inform models aiming to predict species range expansion limits (Dimond et al., 2013), or they can be used to isolate the genomic determinants of thermal tolerance (Dixon et al., 2015; Fuller et al., 2020). Signs of local adaptation can follow predictable environmental gradients, even in marine organisms with high dispersal potential (Sanford & Kelly, 2011). In addition, due to the highly predictable nature of temperature differences along latitudinal gradients, these clines may be key for identifying ecologically-relevant loci for thermal adaptation (Schmidt et al., 2008). However, it is important to note that thermally adaptive loci may lead to fixed phenotypes or plastic phenotypes with underlying thermally adapted gene expression (Ehrenreich & Pfennig, 2016). Phenotypic plasticity also enables organisms of the same genotype to occupy divergent thermal regimes (Yampolsky et al., 2014), and this trait is thought to be particularly relevant in intertidal species with wide spatial ranges that experience high environmental variation within their habitats and across their range. These latitudinal gradients may therefore provide important insights into the ecological and evolutionary processes shaping intertidal species distributions.

Many studies of population structure in marine invertebrates have historically used a small number of markers, such as microsatellites or mitochondrial DNA (Costantini et al., 2018). While these loci can inform coarse scale relationships between temperature and genetic structure (Fullard et al., 2000; Hübner et al., 2009), using a limited number of markers may provide an incomplete picture of more fine scale patterns. Previous work has often detected marine invertebrate population structure across wide spatial scales of hundreds to thousands of kilometers (for review see: Taylor & Roterman, 2017), but the low resolution provided by only a few loci makes it challenging to relate this variation to environmental parameters. For example, Junge *et al* (2011) found temporally inconsistent evidence for structure in a spring-spawning salmonid using 19 loci, despite strong physiological evidence for local adaptation to different temperature regimes in several life-history traits. This challenge may be amplified in species with high dispersal capabilities and potentially high gene flow (as in the case for many marine invertebrates), as this can restrict the signal of adaptive loci to just a few important regions. Genomic studies leveraging more modern technologies can provide finer resolution (D’Aloia et al., 2020), allowing for more direct assessments of the relationship between environmental and genomic variation.

Much like how latitude can be leveraged to compare thermal regimes, comparing congeners offers the opportunity to understand conserved mechanisms involved in response to environments. For example, a comparison of congeneric porcelain crabs across a thermal gradient established the role of different morphological and biochemical features that lead to thermal tolerance (Stillman & Tagmount, 2009). Furthermore, contrasting congeners can elucidate links between thermal tolerance and species distributions. For example, by comparing two invasive and two non-invasive beach grasses, it was found that the two species with broad distributions and high invasion success exhibited wider thermal breadths than the other two species with more narrow ranges and low invasion success (Rehage et al., 2020). Finally, population genetic comparisons of congeners offer important insights into how genetic variation and connectivity are shaped by seascapes (Davies et al., 2015) and because closely related species often share similar substitution and mutation rates (Martin & Palumbi, 1993), this allows for more direct comparisons of gene flow.

Two congener species of *Crepidula* (*C. fornicata*, *C. plana*) live in sympatry along the northeast Atlantic coast of the United States and offer a promising comparative system to characterize how temperature shapes genetic variation through neutral and selective forces. These sister species of benthic marine gastropods are commonly known as ‘slipper snails’ and have a protandric hermaphrodite lifestyle with smaller males forming gregarious stacks on top of larger females (Dupont et al., 2006). Recent attention has focused on *C. fornicata* as an emerging model system for Lophotrochozoan development (Henry et al., 2010), as well its use in CRISPR/Cas-9 mediated gene knockouts (Perry & Henry, 2015). While a promising model system, *C. fornicata* has received most of its attention due to its invasive success in Europe and on the west coast of North America (Blanchard, 1997), which represents a key difference between its native congener. Invasive success in *C. plana* is limited, despite sharing similar life history characteristics and recorded opportunities for invasion in Europe (Minchin et al., 1995). While *C. plana* was originally thought to have the same spatial distribution as *C. fornicata* (Hoagland, 1977), sequencing has uncovered three distinct cryptic species within the *C. plana* complex: *C. plana*, *C. depressa* and *C. atrasolea* (Collin, 2001). These cryptic species of *C. plana* separate between known biogeographic breaks where current dynamics form natural dispersal barriers for many marine organisms (Pappalardo et al., 2015) and hereafter any reference to *C. plana* refers to the species within the complex, not the complex itself. It is possible that *C. plana*’s pelagic larval duration (PLD) contributes to its truncated range given that its maximum PLD is lower than *C. fornicata*’s (12 days vs 21 days; Lima & Pechenik, 1985). The duration of an organism’s PLD can also contribute to genetic structure, where shorter PLD leads to greater differentiation (Bohonak, 1999). However, a meta-analysis on reef invertebrate connectivity found little support for PLD correlating with genetic structure (Costantini et al., 2018). It remains unclear whether PLD differences between congeneric *Crepidula* explain variation in range sizes or whether other biotic or abiotic factors explain these differences.

Here we compare how genetic variation between the congeneric gastropods *C. fornicata* and *C. plana* is partitioned across five sites along the northern portion of their native range in the United States using reduced representation DNA sequencing (2bRAD-seq). We then explored how temperature variation may explain patterns of genetic diversity across this latitudinal gradient. Our findings contribute to growing research exploring how different evolutionary processes drive genetic variation in marine organisms across latitudinal gradients.

## Methods

### Sample collection and environmental data

During the low tides of July and August 2021, individuals identified as *Crepidula fornicata* and *C. plana* were collected from five different sites along the Northeast shores of the United States (Figure 1A). Animals were removed from intertidal substrate using a flathead screwdriver and transported in a cooler containing continuously aerated sea water. No *C. plana* were found at the northernmost sampling site (Robbinston, ME). All animals were transferred into holding aquaria at Boston University’s Marine Invertebrate Research Facility within 24 hours of collection, before being removed from their shells with tweezers, preserved in 200 proof ethanol and stored at −80°C until processing. To characterize seawater and air temperature profiles across sites, hourly temperature data between January 2020 to January 2021 were obtained from the National Oceanic and Atmospheric Administration (NOAA) weather buoys that were closest to each sample location (Table S2). These temperature data were plotted along with a local regression line (loess smoothing) by site location to illustrate seasonal variation in temperature across sites (Figure 1B-C). Average temperatures for each site were calculated for spring (March, April, May), summer (June, July, August), fall (September, October, November), and winter (December, January, February) and these values were used for downstream association analyses.

**Figure 1.**
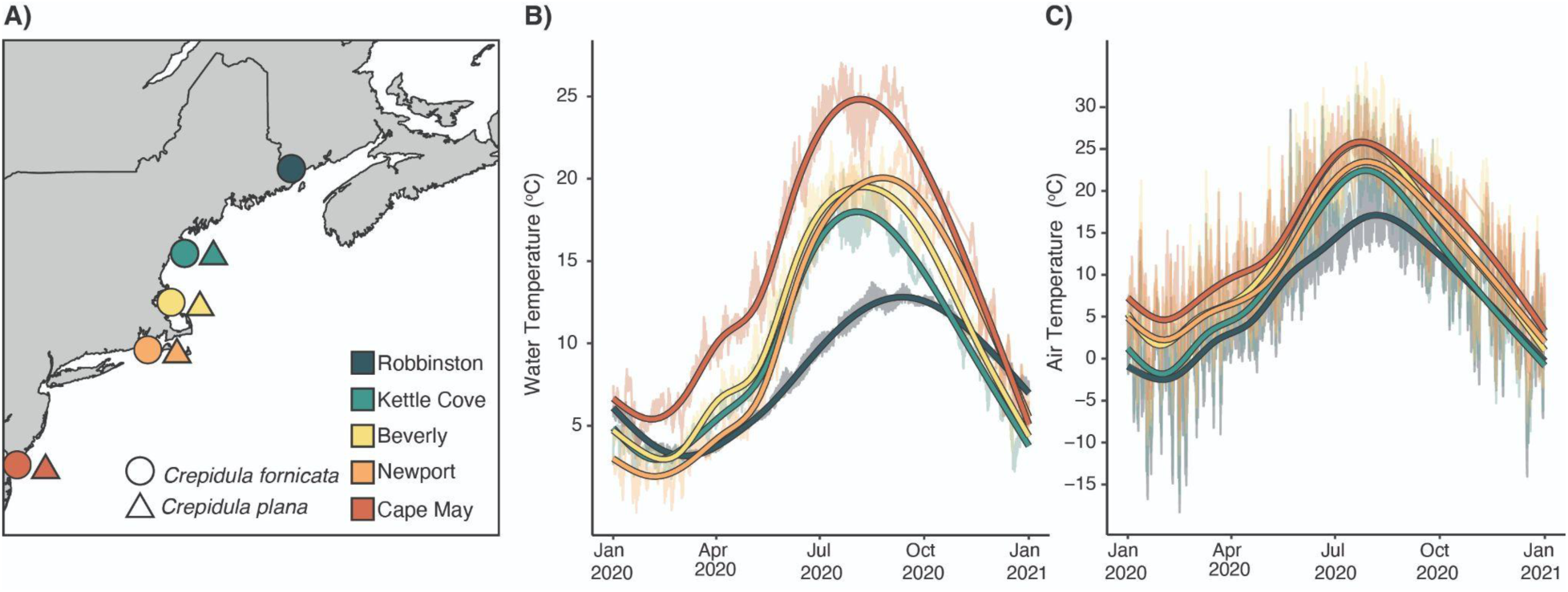
Collection sites and associated temperature profiles for *Crepidula fornicata* and *C. plana*. A) Sampling locations of *Crepidula spp*. along the northeast shore of the United States. Mean hourly temperature plotted along with bolded local regression line (loess smoothing) collected from NOAA weather buoys (Table A3-1) from January 2020 - January 2021 for B) sea surface temperature, and C) air temperature.

### 2bRADseq Library preparation

A small portion of the animal’s foot (approximately the size of a rice grain) was removed with a scalpel and DNA was isolated using Omega BioTek EZNA Mollusc DNA kit following manufacturer’s instructions with one additional DNA washing step (three washing steps total). DNA was normalized to 100 ng in 4uL and prepared for 2b-RAD sequencing following (Wang et al., 2009). Six samples were prepared in duplicate to serve as technical replicates to assist with downstream bioinformatic analyses. A total of 173 animals (see Table S1 for sample sizes across species and sites) were barcoded and sequenced across three lanes of Illumina HiSeq 2500 using single end 50 bp at Tufts University Core Facility.

### Filtering, quality control and single nucleotide polymorphism identification

Per-sample library sizes and *de novo* mapping rates are available in supplemental Table S2. Raw reads were deduplicated, trimmed and quality filtered using *FASTX toolkit* (http://hannonlab.cshl.edu/fastx_toolkit) with Phred scores > 15 (-q 15,15 -m 36). Since no reference genomes are publicly available for either *C. plana* or *C. fornicata, de novo* references were created for each species separately. To construct these references, common tags (major alleles) were collected with the minInd filter set as 10% of sample numbers and then any tag having more than seven observations without a reverse-complement was discarded. References were made using *bowtie2* (Langmead & Salzberg, 2012) setting the chromosome number to 17, based on known chromosome numbers for a related species *Crepidula unguiformis* (Libertini et al., 2009). Sequence alignment files (SAM) from all samples, including technical replicates, were then compressed into BAM files using *samtools* (version 1.12) before inferring initial genotype likelihoods and identification of single nucleotide polymorphisms (SNPs) using *angsd* (version 0.935; Korneliussen et al., 2014) for filters used see Table S3. Seven of 173 samples (4% of samples; Table S4) had >85% missing data and were subsequently discarded. *Angsd* was re-run after these low-coverage samples were removed (for filters see Table S4). The clustering of technical replicates was verified on a dendrogram with hierarchical clustering using the identity by state matrix output from *angsd* using *hclust* in R (version 4.0.2; Figure S1). All technical replicates showed strong clustering (< 0.2 height) in both *C. fornicata* and *C. plana* independent references with initial *angsd* minInd filter set to 80% of respective sample sizes.

### Identifying loci under selection

Candidate loci that are putatively differentiated within a species due to selection were determined using *pcadapt* (Luu et al., 2017). This method identifies candidate SNPs as outliers with respect to population structure. This structure was therefore first inferred using a principal component analysis (PCA) using *PCAngsd* (version 1.11; Meisner & Albrechtsen, 2018), which estimated a covariance matrix of individual allele frequencies from the genotype likelihood files generated by *angsd* described above. *PCAngsd* also generated principal component selection statistics, which were used by *pcadapt* to perform a genome-wide selection scan to identify outlier SNPs. Outliers for each dataset were determined by the Mahalanobis distance of z-scores along the first principal component followed by a ꭓ^2^ test with the outlier SNP alpha set as p < 0.001 (i.e., significantly different from mean distance for each locus in PCA space). Neutral SNPs were identified using this same method, but with a SNP alpha set to p > 0.05, thus some SNPs were not considered in either neutral or outlier analyses. The lists of outlier and neutral SNPs for all datasets were then separately re-run in *angsd* using the same filters as above to determine how genetic variance is shaped when exploring i) all loci, ii) neutral loci, or iii) loci under selection.

### Assessing population structure, genetic diversity and demographic history

Within each species a multidimensional scaling (MDS) analysis was used plotting the Euclidean distances between all samples. This analysis was performed on a covariance matrix for i) all loci, ii) neutral loci and iii) outlier loci based on single-read resampling calculated in *angsd* described above. The position of samples along the first and second MDS axes were visualized, and a permutation multivariate analysis of variance determined significant differences in clustering based on sample site and species using the adonis function in *vegan* (Oksanen et al., 2019). Admixture between individuals within a species was explored using *NGSadmix* (Skotte et al., 2013), which analyzes genotype likelihood files from all SNPs and neutral SNPs only. Optimal K values were determined using the Evanno method (Evanno et al., 2005) by bootstrapping the above analyses 10 times for each K value between 2 and 5, and using the log likelihood of the estimates from each iteration as input into *CLUMPAK* (Kopelman et al., 2015). To determine genetic differentiation within each species between collection sites, *angsd* calculated site allele frequencies (SAFs) for each population. Then, *realSFS* calculated the folded site frequency spectrums (SFSs) for all possible pairwise comparisons. These folded SFSs were used to calculate Global Watterson’s θ, π, and Tajima’s D for each population using the *thetaStat* tool in *angsd*. Mean differences in these metrics among sampling locations and species were assessed using a Dunn’s test with multiple test correction using the Bonferroni method (Hochberg, 1988). SFSs were then used as priors with the SAF to calculate pairwise F_ST_. Only weighted pairwise F_ST_ values between populations within a species are reported here. To test for patterns of isolation by distance (IBD), a matrix of geographic distances (shortest straight line) between sites was correlated with a matrix of pairwise F_ST_ values using a Mantel’s test with all possible permutations implemented in *vegan*. Finally, to reconstruct the history of effective population size changes on our folded SFS, *Stairway Plot* v2 (Liu & Fu, 2020) was used, which is an unsupervised method requiring no prespecificied demographic models. For all demographic analyses we used default parameters for mutation rate (1.2e-8 per base per year), and a generation time of 3 years.

### Identifying loci associated with temperature regimes

Ordinances from the MDS analyses for each species were correlated with the average seasonal temperature data obtained from NOAA weather buoys (see Table S1). Locations on the first axis of each MDS, which explained the most genetic variance, were plotted against temperature regimes using linear regressions. A series of redundancy analyses (RDAs) were then used to model the relationship between genomic variation with that of environmental and landscape predictors. These were performed using genotype matrices for all loci and neutral loci and in cases of missing data the most common genotype at each SNP was assigned. The genotype matrix of all loci was used as the response variable and the neutral genotype matrix was used to assess the contribution of neutral population structure in the RDAs. To assign neutral population structure, we first performed a principal component analysis on neutral loci and then used eigenvalues from the first principal component as a variable in our models. Significant environmental variables were chosen based on a forward selection strategy with the *ordi2step* function in *vegan* (Oksanen et al., 2019) with a significance value of p < 0.01 and 1000 permutations. A full model was chosen where genetic variation was predicted from significant environmental variables whose model from the above step had the lowest Akaike information criteria (AIC). This model then further included latitude and neutral genetic structure and was compared with a pure climate model (significant environmental variables as explanatory and latitude and neutral population structure as conditional), pure latitude model (latitude as explanatory and environmental and neutral population structure as conditional), and a pure neutral population structure model (neutral population structure as explanatory with environmental and latitude as conditional) using partial RDAs.

## Results

Across all samples, sequencing resulted in an average of 693,161 (SE=28,787) reads per individual, which was reduced to 617,260 (SE = 26,182) after filtering and mapping to species specific *de novo* references. Within species, mapping of *C. fornicata* (reference: 14.16 Mbp) identified 4,379 total SNPs (outliers: 225; neutral: 3,570). *C. plana* (reference: 15.81 Mbp) had approximately 4.5x as many SNPs (outliers: 317; neutral:17,322).

### Population Structure

Multidimensional scaling (MDS) plots demonstrate significant population structure across sites for *C. fornicata* regardless of the loci being tested (All = 4,379, Neutral = 3,570, Outlier = 225) (Figure 2; B,E,H, p_sites_ < 0.001). In contrast, *C. plana* exhibited significant population genetic structure for all loci (p_sites_ < 0.001) and outlier loci (p_sites_ < 0.001), but no structure was observed for neutral loci (p_sites_ = 0.771) (Figure 2; C, I, F;). Population structure was further explored using all loci and neutral loci using ADMIXTURE, which identified (K = 3) ancestral populations in both *C. fornicata* (Figure S2) and *C. plana* (Figure S3). These admixture analyses showcased ancestral population assignments shifting along the latitudinal gradient (Figure S2, S3). When pairwise genetic divergences were calculated between sites (Table S5; F_ST_), all comparisons were significant, but low in magnitude (<0.018), with the largest F_ST_ being between sites with the greatest distance in both species. This pattern resulted in significant isolation by distance (IBD) for *C. fornicata* (Figure 3; Mantel’s r = 0.760, p = 0.0167), but not for *C. plana* (Mantel’s r = 0.658, p = 0.083). However, this IBD signal in *C. fornicata* was no longer significant when the most northern sampling site (Robbinston, ME) was removed from the *C. fornicata* analysis (r = 0.627, p = 0.083) and both the slope and correlation-coefficient were similar in both species, suggesting that the *C. plana* dataset was potentially underpowered.

**Figure 2.**
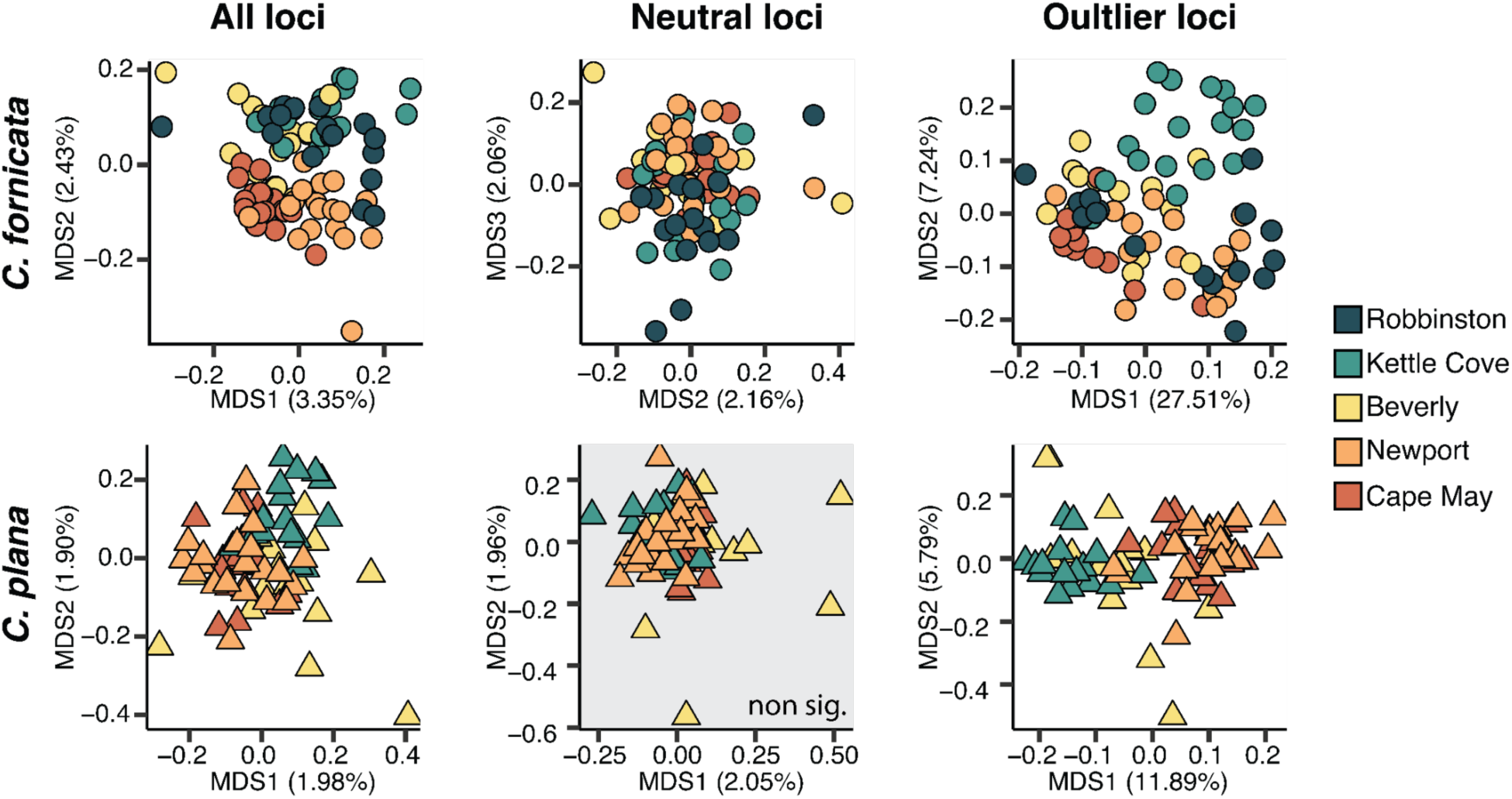
Multidimensional scaling (MDS) plots showing population genetic structure of *Crepidula fornicata* and *C. plana*. Structuring is shown for single nucleotide polymorphisms (SNPs) across all loci, and those that were identified as neutral (p > 0.05), and outlier (p < 0.001) SNPs via *pcadapt*. Significant genetic structure was detected across sampling sites using a permutational multivariate analysis of variance in all analyses except for neutral loci in *C. plana* (denoted with grey box).

**Figure 3.**
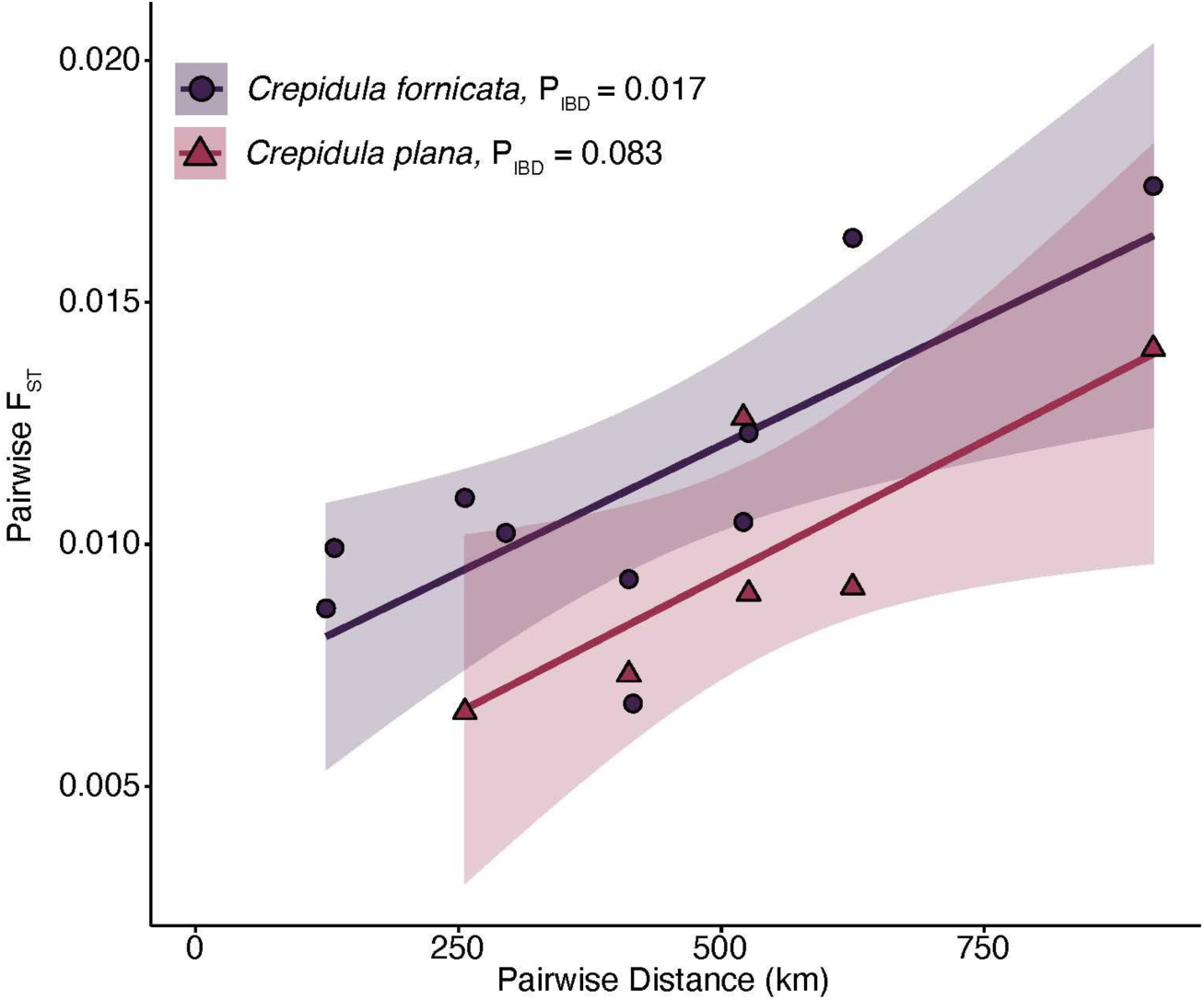
Estimated isolation by distance (IBD) as measured by the relationship between pairwise physical distances between sites (km) and pairwise genetic divergences (F_ST_) in *Crepidula fornicata* (purple) and *C. plana* (pink). IBD was calculated using a Mantel’s test. *C. fornicata* exhibited significant IBD, but *C. plana* did not, potentially due to its relatively lower power (see text).

### Genetic Diversity

Global Watterson’s θ was greater in *C. plana* (Kruskal-Wallis ꭓ^2^ = 5.54, df = 1, p = 0.0186); however, no difference in genetic diversity (π) between species was observed (Kruskal-Wallis ꭓ^2^ = 2.42, df = 1, p = 0.120). *C. fornicata* had a greater average Tajima’s D across all populations (Kruskal-Wallis ꭓ^2^ = 14.0, df = 1, p < 0.001). Comparisons of the sampling locations within species show no differences in π between locations in both *C. fornicata* and *C. plana*. Within *C. fornicata,* Watterson’s θ was significantly lower and Tajima’s D higher in Robbinston, ME compared with Newport, RI and Cape May, NJ (Figure 4B-C). Within *C. plana,* Watterson’s θ was significantly lower and Tajima’s D higher in Beverly, ME compared to all other sampling locations (Figure 4B-C).

**Figure 4.**
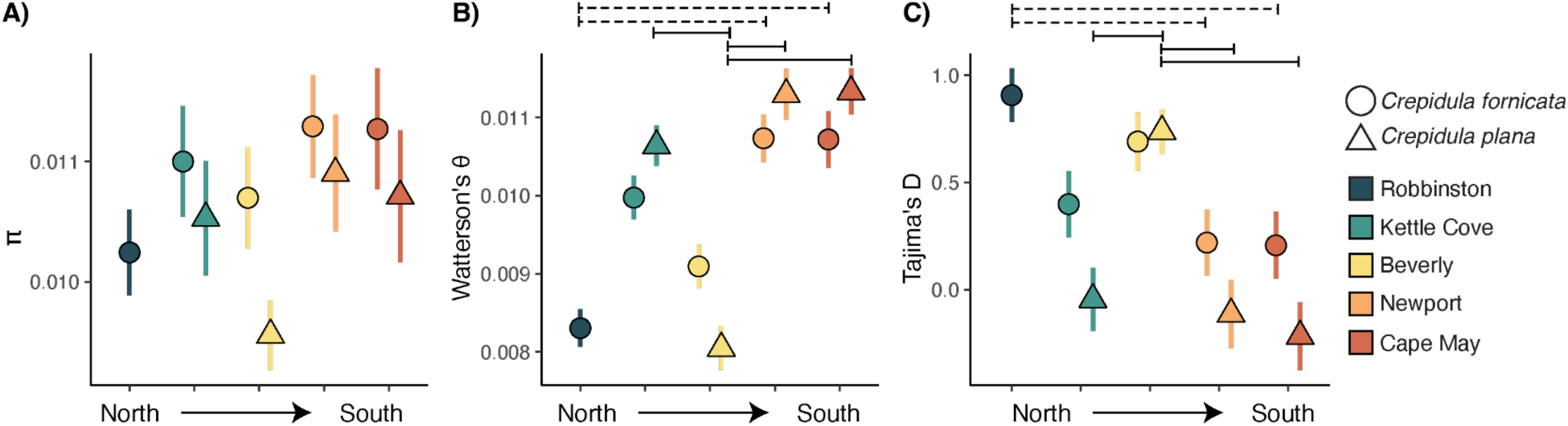
Genetic diversity estimates. A) genetic diversity (π), B) Watterson’s θ, and C) Tajima’s D. Points represent means, with error bars denoting standard error. Significant pairwise comparisons (p_adj_ < 0.05) within species are denoted by dashed horizontal lines for *Crepidula fornicata* and solid lines for *C. plana*.

### Demographic inference

Stairway plot analysis of effective population (N_e_) sizes of *C. fornicata* suggest recent N_e_ contractions over the past few thousand generations at all sites except for Newport, RI (Figure 5A). These contractions vary in their magnitude and follow the latitudinal gradient with Robbinston, ME having the smallest contemporary N_e_ and increasing at each site moving southward. This contrasts with *C. plana’s* results, which suggest increasing N_e_ over 100,000 generations ago followed by relative stasis, except for Beverly, MA (Figure 5B). In this location, the trend follows a similar pattern to *C. fornicata* with contemporary contractions in N_e_.

**Figure 5.**
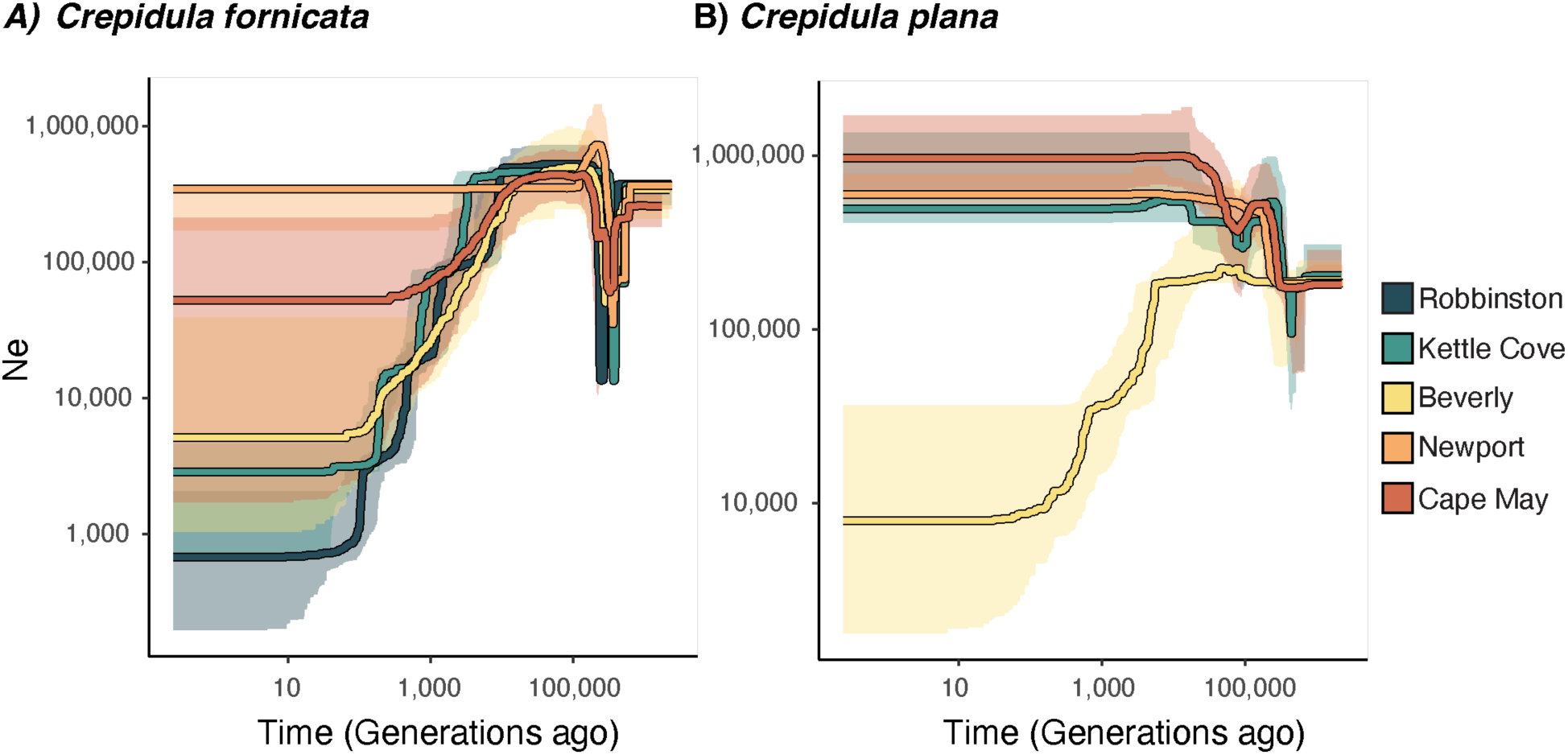
Stairway plot of inferred effective population (N_e_) sizes over time for A) *Crepidula fornicata* and B) *C. plana*. Lines show median N_e_ and ribbons represent 95% confidence intervals. Note that y-axis scales differ between species.

### Associations of temperature and genetic variance

Two approaches were used to measure the association of temperature with genetic variation. The first approach associated ordinations along the first MDS axis for i) neutral, and ii) outlier loci with average seasonal (spring, summer, winter, fall) air and water temperatures. Using this method, structure in neutral loci was correlated with seasonal temperatures in *C. fornicata* (p < 0.05), except for winter water temperatures not showing a significant association (Figure S4). This pattern contrasted with *C. plana* where structure in neutral loci exhibited no associations with seasonal water and air temperatures, except for summer and spring air temperatures exhibiting weak associations (Figure S5). Lastly, structure in outlier loci showcased the strongest associations with water and air temperatures in both species. In *C. fornicata*, structure observed in outlier loci was correlated with temperatures across seasons (Figure S6; all p < 0.001) and similar patterns were observed in *C. plana* except for spring sea surface temperature having no significant association (Figure S7).

While the above approach assesses correlations between genetic variation and temperature, we also employed a multivariate approach to further characterize how temperature and spatial heterogeneity impact genetic variation using a series of redundancy analyses (RDAs; Figure 6). When using a forward selection strategy to select a pure temperature model that best explained our data, we found that summer sea surface temperature in *C. fornicata* and fall sea surface temperatures in *C. plana* were the best fit models (lowest AIC). When these top environmental models were used as the explanatory variables, while controlling for latitude and neutral population structure in an RDA, temperature contributed 9% of the explainable variance in *C. fornicata* and 12% in *C. plana*. Latitude was also a significant predictor of the explainable variance in both species (10% *C. fornicata*, 12% *C. plana*) when controlling for temperature and neutral population structure. Neutral population structure explained the most variation in both species (*C. fornicata*: p < 0.05, 77%; *C. plana*: p < 0.05, 71%), when controlling for latitude.

**Figure 6.**
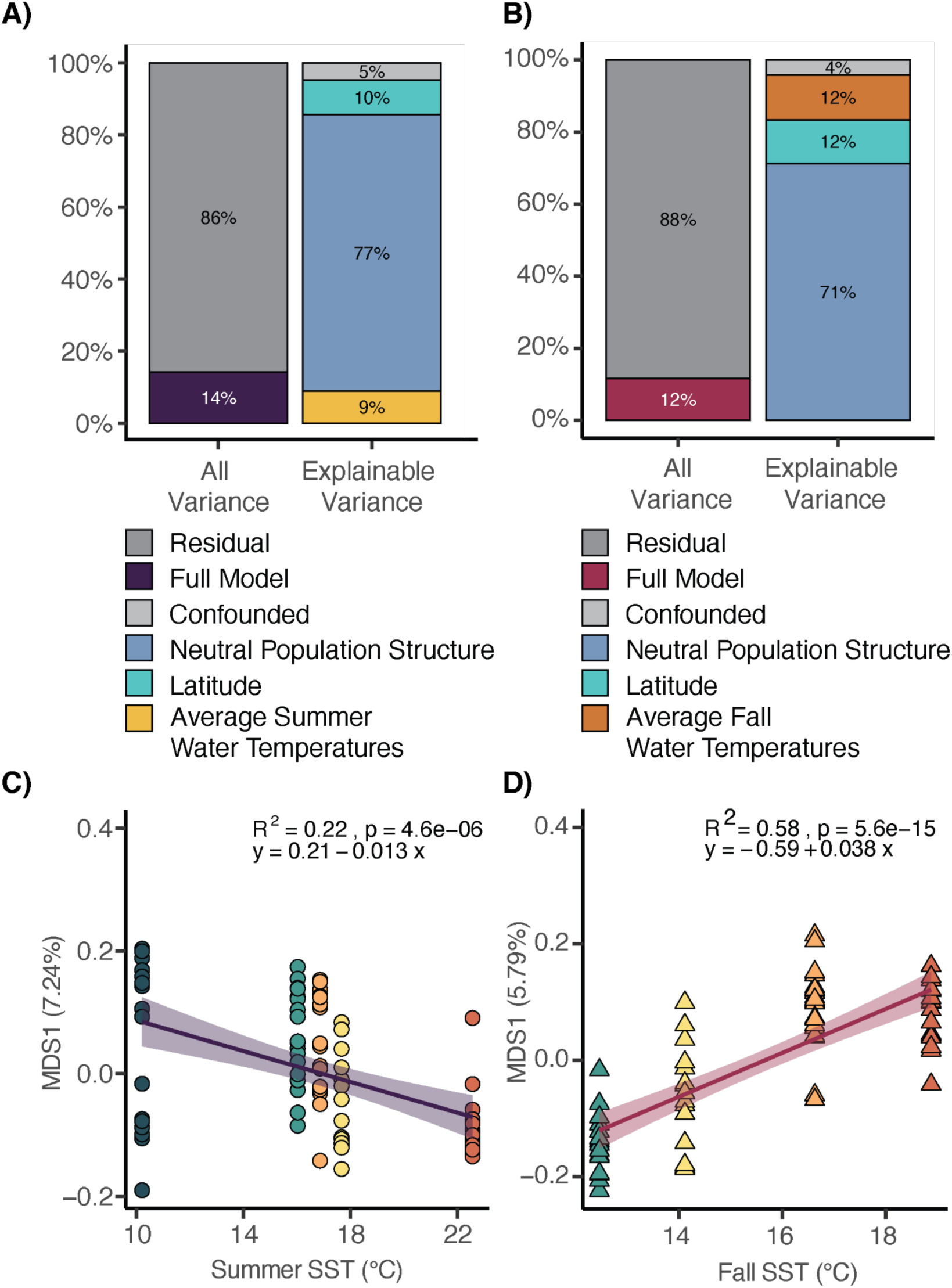
Environmental variation associated with genetic variation in all loci of A) *Crepidula fornicata* and B) *C. plana*. Bar plot denotes variance explained in redundancy analysis (RDA). Confounded percentage denotes variation that could not be separated between temperature models (summer temperature in *C. fornicata* and fall temperature in *C. plana*) and geographic models (latitude). C) Average summer water temperatures were a significant driver of variation for *C. fornicata* and were correlated with genetic variation of outlier loci using the first MDS axis from Figure 3. D) Average fall water temperatures were a significant driver for *C. plana* genetic variation within outlier loci using the first MDS axis from Figure 3.

## Discussion

We investigated patterns of genetic variation in the two congeneric marine gastropods *Crepidula fornicata* and *C. plana* across a latitudinal gradient (approximately 900 km) representing a significant temperature cline spanning the northern portion of their native ranges. These species share many similar life history traits such as planktonic larval development yet exhibit key differences in the extent of their spatial ranges. *C. fornicata* exists across a more extensive local range than *C. plana* (Collin, 2001) in addition to having additional contemporary invasive ranges globally (Blanchard, 1997). We inferred demographic histories across this latitudinal gradient and tested for associations between genetic variation and temperature. Broadly, similar divergence patterns were observed between *C. fornicata* and *C. plana* across latitude, with *C. fornicata* exhibiting higher genetic divergence between locations. Our demographic analyses revealed that contemporary contractions of effective population (N_e_) sizes in *C. fornicata* followed a latitudinal gradient where N_e_ contractions were higher at higher latitude sites. In contrast, *C. plana* population sizes have remained much more constant. Finally, temperature regimes shaped genetic variation in both neutral and outlier loci in *C. fornicata*, but only outlier loci in *C. plana*. These results suggest subtle differences in genetic differentiation between these congeners, perhaps due to differences in gene flow between populations or historic bottlenecks in *C. fornicata* that were not experienced by *C. plana.* Our results support a possible role for temperature in shaping divergence patterns across the seascape, though further work disentangling temperature from other abiotic factors that are correlated with latitude is needed.

We detected significant genome-wide population structure across the latitudinal gradient in all, putatively neutral, and selected loci in *C. fornicata* (Figure 2). Similar patterns emerged in *C. plana* in all and selected loci; however, there was no significant structuring in neutral loci (Figure 2). These patterns were consistent with our admixture results, where a distinct north to south gradient in population structure for all loci and neutral loci was observed in *C. fornicata* (Figure S2) whereas *C. plana* only exhibited latitudinal associations in all loci and not neutral loci (Figure S3). While population structure was observed, low between-site F_ST_ was also found in both species, suggesting high gene flow across this range (Table S5). We found a significant pattern of isolation by distance (IBD; Kimura & Weiss, 1964), in *C. fornicata* but not in *C. plana* (Figure 3); however both species exhibit similar IBD characteristics, and this trend could potentially become significant with additional sites sampled. While different structure patterns between species could be driven by a discrepancy of the total number of loci, we do not think this is the case here as we found similar numbers of total loci in the *de novo* references in both species. Overall, each of our analyses paints a consistent picture that population structure along the latitudinal gradient exists but is relatively weaker in *C. plana*.

These findings are largely consistent with previous work in *Crepidula,* which have explored population structure using few loci. For example, genetic structure in *C. fornicata* but not *C. plana* was found along the coast of New England using cytochrome oxidase I (Collin, 2001) and allozymes (Hoagland, 1985). Our *C. fornicata* results are also similar to those observed using microsatellite markers across their native range (Riquet et al., 2013; Viard et al., 2006). Together, our results provide further evidence that biophysical features of *C. fornicata* across its native range lead to genetic divergence across the seascape, and these same features impact *C. plana,* although to a lesser degree. Work in other intertidal invertebrates such as in mussels (*Mytilus edulis*; Riginos & Henzler, 2008), oysters (*Crassostrea virginicacite*; Bernatchez et al., 2019), and barnacles (*Pollicipes pollicipes*; Quinteiro et al., 2007) have all consistently shown evidence for neutral population structure, highlighting that the apparent high connectivity of *C. plana* may make it an outlier amongst invertebrates in this region.

In addition to population structure, we also measured differences in genetic diversity as well as inferred demographic histories across the latitudinal gradient in both species. In *C. fornicata* we found lower Watterson’s θ and higher Tajima’s D in samples from the most northern sampling site relative to two southern locations (Newport, RI, Cape May, NJ; Figure 4B-C). This pattern suggests a depletion of rare alleles in the north and may correspond with inferred decreases in N_e_ over the last few thousand years (Figure 5A). This recent population contraction is contrary to demographic signals inferred in *C. plana* as well as other temperate marine invertebrates such as scallops whose northern populations show steady N_e_ expansions over the last few thousand years (Vendrami et al., 2019). *C. plana* maintained consistently high N_e_ at all sites except Beverly, MA, which had a much smaller contemporary N_e_ and demographic history that more closely mirrors pattern observed in *C. fornicata*. This site is a clear outlier in both diversity statistics as well as in our demographic analysis. Our demographic analyses are consistent with a model in which *C. plana* populations have remained resident to the northeast, while *C. fornicata* populations either declined during the last glacial maximum or reinvaded as glaciers receded. This process of population contraction may have increased signals of IBD in *C. fornicata* due to founder effects not experienced by *C. plana*. In this model, the northern populations represent the leading edge of an expanding wavefront, perhaps beginning at the end of the last glacial maximum, rather than a contracting native population. Future research using annotated whole genome sequences and haplotype patterns may further clarify the specific drivers of differentiation in these populations over their evolutionary histories.

There are many plausible explanations for the differences observed here between *C. fornicata* and *C. plana*. *C. plana* was not observed at the northernmost site, perhaps because it has a smaller thermal niche width than *C. fornicata*. A more truncated niche space could lead to patterns of population structure in *C. plana* to become less pronounced simply because it has a smaller geographic range. Another key difference between these species is that *C. plana* tends not to form large, gregarious stacks like *C. fornicata*. These stacks function as independent mating groups, often with skewed sex ratios with more males than females (Collin, 1995). Therefore, despite a larger census population size, *C. fornicata’s* stacking behaviour could lower their N_e_. This would increase neutral genetic divergence between populations as this stacking behavior could increase inbreeding potential, and the influence of genetic drift (Wright, 1980). While this is certainly the case in gonochoric organisms (Frankham, 1995), hermaphroditic organisms may be able to overcome these disadvantages as individuals can breed as both sexes. For example, work in fishes has demonstrated that sequential hermaphroditism is an evolutionary stable strategy that minimizes the evolutionary constraints (such as reduced N_e_) of a skewed sex ratio (Waples et al., 2018). *Crepidula* therefore presents opportunities for future research to understand how these unique reproductive characteristics may impact effective population size, rates of inbreeding, or other aspects of biology that may drive the population genetic patterns observed between *C. fornicata* and *C. plana*.

### Genetic differentiation at putative loci under selection associates with temperature in *C. fornicata* and *C. plana*

Species in intertidal zones experience frequent fluctuations in temperature and environmental conditions due to the changing tides (Lima et al., 2011). These fluctuations become more pronounced at higher latitudes, with both temperature and tidal extremes intensifying. Therefore, adaptations to broad thermal regimes are necessary to survive these harsh conditions (Diederich & Pechenik, 2013). Here, we find support for temperature shaping genetic variation across a latitudinal gradient in both *Crepidula* species, and this pattern was most prominent in putatively selected loci. Correlations between temperature and MDS1 of neutral loci were either non-significant or weakly significant in both *C. fornicata* (Figure S4) and *C. plana* (Figure S5). In contrast, associations using outlier (putatively selected; Figure S6, S7) loci alone were considerably stronger in both species. Therefore, these loci might be driven by adaptation to thermal regimes rather than through neutral processes, though we cannot rule out neutral processes completely (see discussion of RDA below). Previous work by Thieltges et al., (2004) concluded that winter temperatures were the primary constraint for range expansion in northern populations of *C. fornicata*. From their results, we might expect that winter months would impart strong selective pressures and therefore be strongly correlated with outlier loci observed here. It has also been proposed that cold water acts as a significant selective force in *C. plana,* with limitations in cold water tolerance potentially explaining why *C. plana* has a more limited northern range and has failed as an invasive species in Europe (Boisvert, Kelly, 2014). Our results in both analyses fail to support this prediction as the association between temperature and differentiation amongst outlier loci was strongest during the summer (*C. fornicata*; Figure 6A) and fall (*C. plana*; Figure 6B) when water temperatures are the warmest in this region. Together, our data suggest that patterns of adaptive genetic divergence in *Crepidula* are more strongly associated with warmer temperatures. Future research characterizing thermal performance curves of these populations coupled with high resolution whole genome sequencing could provide further support for this hypothesis.

Genomic signatures of selection can sometimes be falsely attributed to variance that is evolutionarily neutral (Excoffier & Heckel, 2006). A limitation of the correlation analyses we performed between outlier loci and temperature variables is that outlier loci are detected because they have larger than expected frequency differences between populations, which means their latitudinal signals are inherently stronger. Therefore, we controlled for neutral population structure by performing a redundancy analysis, which is a more conservative approach to identify signals of selection (Forester et al., 2018). We found that after controlling for latitude and neutral population structure, temperature explained approximately 9% of explainable variance in *C. fornicata* and 12% in *C. plana* (Figure 6). While temperature explained similar amounts of genetic variance in both species, our redundancy analysis highlights that different seasons contribute to this variance between congeners. Summer (*C. fornicata*) and fall (*C. plana*) water temperatures explained the most variation, suggesting that warmer waters are stronger drivers of this genetic variation. This pattern contrasts with many terrestrial species where winter lows have the largest explanatory power (Ashcroft et al., 2011). While this may be true for terrestrial plants, likely due to frost and freezing damage, this aspect of temperature may not be as relevant in marine organisms. Indeed, RDA approaches often identify warm water as explaining adaptive genetic variance in marine invertebrates at latitudes like those sampled here. For example, warmer sea temperatures explained genetic variation in sea cucumbers (*Parastichopus californicus;* Xuereb et al., 2018), and urchins (*Strongylocentrotus purpuratus*; Pespeni & Palumbi, 2013) along latitudinal clines. However, we only found that 9-12% of the explainable variance was associated with temperature. It is possible that more finely tuned temperature data that better matches local conditions of sampling sites would better inform the RDA models. Here, we used oceanic weather buoy data, which likely underestimates temperature variation experienced by these intertidal organisms. Biomimetic “robosnails” temperature loggers placed amongst *C. fornicata* populations in Rhode Island, USA showed that these organisms experience extreme diel temperature variability (20-43°C; Diederich & Pechenik, 2013). Future work would benefit from the use of satellite-based weather data or through incorporating “robosnails” temperature loggers at each site. Nevertheless, despite these limitations we show evidence for temperature explaining genetic variance and this variation is likely adaptive.

## Conclusion

We compared population genetic patterns in two congeneric marine gastropods with similar life history characteristics in the genus *Crepidula* to explore how genetic variation is partitioned across a wide latitudinal gradient. We found parallel patterns of genetic divergence and diversity between species, with subtle differences in connectivity across the seascape. Notably, in contrast to our initial predictions based on range size, *C. fornicata* showed stronger evidence for isolation by distance and higher neutral population divergence compared to *C. plana,* which has a narrower range. However, both species demonstrate adaptive differentiation across the latitudinal gradient, with significant associations with temperature. These findings imply nuanced variation in gene flow within the coastal landscape among these closely related species, with temperature shaping these population genetic patterns. We therefore predict that under future ocean warming associated with climate change, these warmer temperatures will continue to influence genetic variation in these species.

## Acknowledgements

We extend gratitude towards Ashley Phan, Angela Xia, Hanna Kim and Kimberly Candelario for their assistance with DNA extractions. We acknowledge Boston University’s Shared Computing Cluster (SCC) for computational resources associated with these analyses. Members of the Davies and Uricchio lab for their thoughtful feedback and perspectives from Pete Buston, Randi Rotjan, Sean Mullen and Rachel Wright throughout the analysis, interpretation, and editing process.

## Data Availability

Code and files can be accessed: https://github.com/wuitchik/crepidula

