## Supplemental Materials for "Outside your shell: how temperature shapes genetic variation in two species of congeneric marine snails"

**Table S1:** Site locations, sample sizes and closest NOAA weather buoy information.

| Site Name | Site Location | NOAA ID | NOAA Buoy Location | $N_{C. fornicata}$ | $N_{C. plana}$ |
|---|---|---|---|---|---|
| Robbinston, ME | 45°05'01.8"N 67°06'36.1"W | PSBM1 | Eastport, ME | 16 | 0 |
| Kettle Cove, ME | 43°33'40.8"N 70°13'08.3"W | CASM1 | Portland, ME | 18 | 16 |
| Beverly, MA | 42°32'48.2"N 70°51'46.9"W | BHBM3 | Boston, MA | 17 | 19 |
| Newport, RI | 41°34'29.3"N 71°17'13.8"W | NWPR1 | Newport, RI | 19 | 20 |
| Cape May, NJ | 38°58'46.8"N 74°57'40.8"W | CMAN4 | Cape May, NJ | 20 | 20 |

**Table S2:** Read numbers across quality control and filtering steps.

| Sample | Raw | Filtered | Mapped to own species | Percent missing data after initial *angsd* run | Notes |
|---|---|---|---|---|---|
| FB11A | 571064 | 563209 | 542253 | 20.5% | |
| FB12A | 260991 | 257504 | 244927 | 88.3% | Removed for high missing data |
| FB13A | 1138740 | 1124240 | 1047411 | 1.3% | Concatenated with FB13B |
| FB13B | 724072 | 715403 | 685425 | 7.4% | |
| FB14A | 402071 | 396472 | 383897 | 37.8% | |
| FB14B | 361130 | 357151 | 340812 | 12.9% | |
| FB21A | 183971 | 181396 | 159982 | 84.4% | |
| FB21B | 563324 | 554856 | 528108 | 13.5% | |
| FB22A | 562432 | 555277 | 521756 | 31.7% | |

| | | | | | |
|---|---|---|---|---|---|
| FB22B | 901549 | 890420 | 840269 | 8.1% | |
| FB23A | 650314 | 641108 | 597598 | 5.5% | |
| FB23B | 226496 | 223375 | 216380 | 85.3% | Removed for high missing data |
| FB23C | 1020692 | 1007368 | 943590 | 1.3% | |
| FB32A | 554688 | 546345 | 478346 | 64.9% | |
| FB41A | 331329 | 327527 | 311121 | 12.8% | |
| FB42B | 584647 | 577205 | 545325 | 3.9% | |
| FB44A | 1202028 | 1185695 | 1111045 | 0.7% | |
| FC12A | 415777 | 411035 | 388561 | 8.3% | |
| FC12B_2 | 965364 | 954236 | 891085 | 2.5% | Concatenated with FC12A |
| FC12B | 826276 | 816780 | 767334 | 2.2% | |
| FC13A | 1362533 | 1342830 | 1261173 | 0.9% | |
| FC14A | 1607702 | 1586217 | 1478065 | 0.7% | |
| FC21A | 587853 | 579680 | 535254 | 23.6% | |
| FC21B | 313422 | 309724 | 294960 | 14.9% | |
| FC22A | 606841 | 599184 | 551341 | 28.8% | |
| FC22B | 1130669 | 1114164 | 1060792 | 1.5% | |
| FC24A | 759738 | 750762 | 708487 | 3.2% | |
| FC24B | 897031 | 886642 | 830682 | 2.1% | |
| FC31A | 775618 | 766495 | 714038 | 5.4% | |
| FC32A | 761501 | 750486 | 704530 | 3.2% | |
| FC32B | 734462 | 723261 | 689418 | 10.0% | |
| FC33A | 809483 | 799192 | 731979 | 1.4% | |

| | | | | |
|---|---|---|---|---|
| FC33B | 273319 | 270388 | 253428 | 23.6% |
| FC33C | 1425768 | 1407836 | 1319362 | 0.8% |
| FC34A | 421365 | 416666 | 386596 | 17.2% |
| FC41A | 422296 | 417052 | 384336 | 50.1% |
| FC42A | 717302 | 706229 | 655826 | 34.7% |
| FC43A | 629817 | 621579 | 513525 | 53.2% |
| FK11B | 671783 | 663422 | 620796 | 3.6% |
| FK12A | 392264 | 386785 | 366370 | 11.5% |
| FK21A | 247860 | 244698 | 232969 | 23.8% |
| FK21B | 177352 | 174688 | 161539 | 45.3% |
| FK22A | 1007285 | 993996 | 909354 | 2.4% |
| FK23A | 759751 | 749490 | 706564 | 2.5% |
| FK24B | 423505 | 418412 | 394181 | 12.8% |
| FK31A | 1295775 | 1278536 | 1181827 | 1.4% |
| FK33A | 674717 | 667287 | 619983 | 6.3% |
| FK33B | 219390 | 216940 | 203531 | 40.8% |
| FK33C | 167712 | 165661 | 154774 | 59.4% |
| FK34A | 1099018 | 1083404 | 1008989 | 0.8% |
| FK34B | 265583 | 262593 | 248952 | 22.5% |
| FK41A | 494348 | 487723 | 451996 | 13.4% |
| FK43A | 517693 | 511872 | 477915 | 16.3% |
| FK44A | 469276 | 463847 | 426718 | 7.3% |
| FK44B | 697968 | 690385 | 646586 | 3.9% |
| FK44C | 940363 | 928199 | 851366 | 4.6% |

| | | | | | |
|---|---|---|---|---|---|
| FN11A | 336818 | 332518 | 319542 | 48.6% | |
| FN11B | 258299 | 255357 | 243244 | 18.7% | |
| FN13A | 566659 | 558826 | 467395 | 65.5% | |
| FN14A | 401524 | 396732 | 372977 | 10.1% | |
| FN21A | 972267 | 959350 | 903899 | 1.4% | |
| FN22A | 882631 | 871243 | 810724 | 2.2% | |
| FN22B | 590153 | 581768 | 548724 | 4.2% | |
| FN22C | 858492 | 847435 | 795526 | 2.3% | |
| FN31A | 517920 | 510649 | 481175 | 55.1% | |
| FN32A | 667099 | 659273 | 622327 | 6.8% | |
| FN33A | 296204 | 291263 | 272333 | 96.8% | Removed for high missing data |
| FN33B | 479607 | 474307 | 448362 | 6.0% | |
| FN34A | 803616 | 794085 | 742341 | 6.8% | |
| FN34B | 503061 | 495530 | 466333 | 71.1% | |
| FN43A | 864367 | 853401 | 800858 | 5.5% | |
| FN43B | 1241783 | 1227112 | 1149969 | 0.8% | |
| FN43C | 621830 | 613819 | 579680 | 6.7% | |
| FN44A | 1149495 | 1135726 | 1070776 | 1.2% | |
| FN44B_2 | 296159 | 292444 | 275595 | 19.5% | Concatenated with FN44A |
| FN44B | 490716 | 484431 | 455015 | 7.0% | |
| FN44C | 425635 | 421068 | 396761 | 9.5% | |
| FR13A | 206346 | 203854 | 189119 | 38.4% | |
| FR21A | 1243584 | 1227172 | 1155420 | 0.6% | |

| | | | | | |
|---|---|---|---|---|---|
| FR21B | 1271370 | 1255324 | 1175109 | 0.9% | |
| FR23B | 337768 | 333225 | 314871 | 14.1% | |
| FR23C_2 | 490830 | 485057 | 465528 | 12.9% | Concatenated with FR23_C2 |
| FR23C | 341571 | 337079 | 324282 | 21.0% | |
| FR24A | 714064 | 704860 | 619147 | 13.9% | |
| FR24B | 473720 | 467931 | 436763 | 15.9% | |
| FR31A | 297523 | 293625 | 219912 | 75.4% | |
| FR31B | 245995 | 242481 | 234054 | 69.8% | |
| FR32A | 276467 | 272568 | 258653 | 20.3% | |
| FR331A | 939746 | 927666 | 871503 | 2.4% | |
| FR33A | 174447 | 172077 | 166263 | 79.9% | |
| FR411A | 357109 | 352242 | 333839 | 12.1% | |
| FR43A | 219592 | 216935 | 209143 | 36.8% | |
| FR43B | 154866 | 153022 | 143168 | 37.8% | |
| FR43C | 190994 | 188935 | 180552 | 30.5% | |
| PB11A | 325625 | 320767 | 300595 | 96.1% | Removed for high missing data |
| PB11B | 780713 | 770446 | 230877 | 56.3% | |
| PB11C | 941751 | 930095 | 869143 | 5.7% | |
| PB12A | 537658 | 530722 | 503944 | 15.7% | |
| PB13A | 416658 | 410912 | 387114 | 16.9% | |
| PB13B | 248173 | 244578 | 231403 | 97.3% | Removed for high missing data |
| PB14A | 889194 | 878320 | 825553 | 6.3% | |

| | | | | | |
|---|---|---|---|---|---|
| PB14B | 330811 | 326237 | 310509 | 73.0% | |
| PB21A | 446496 | 439828 | 332848 | 63.4% | |
| PB22A_2 | 725996 | 716878 | 679092 | 8.3% | Concatenated with PB22A |
| PB22A | 642821 | 634262 | 602179 | 13.1% | |
| PB23A | 183638 | 181385 | 170877 | 60.1% | |
| PB24A | 560572 | 553305 | 518843 | 9.3% | |
| PB24B | 644107 | 635303 | 544040 | 8.7% | |
| PB31A | 687410 | 678443 | 601883 | 21.6% | |
| PB32A | 623290 | 615230 | 578802 | 10.0% | |
| PB33A | 218024 | 215100 | 185529 | 65.9% | |
| PB33B | 1536909 | 1517845 | 1421159 | 1.1% | |
| PB33C | 889511 | 876380 | 799489 | 3.5% | |
| PB34A | 447169 | 441013 | 410274 | 36.1% | |
| PB43A | 495889 | 490068 | 463784 | 8.4% | |
| PC11A | 648844 | 640431 | 585037 | 9.3% | |
| PC12A | 1195836 | 1179961 | 1105281 | 2.3% | |
| PC13A | 882417 | 871172 | 808201 | 3.0% | |
| PC13C | 840461 | 830905 | 777325 | 4.1% | |
| PC14A | 1341255 | 1324681 | 1238598 | 2.2% | |
| PC21A | 1445757 | 1426091 | 1335941 | 1.4% | |
| PC22A | 958445 | 944616 | 880786 | 3.3% | |
| PC22B | 271935 | 268186 | 233853 | 87.5% | Removed for high missing data |
| PC22C | 1100931 | 1087304 | 954232 | 2.1% | |

| | | | | | |
|---|---|---|---|---|---|
| PC23A | 498895 | 491430 | 336272 | 16.7% | |
| PC23B | 741075 | 731896 | 683663 | 7.3% | |
| PC23C | 1101891 | 1088166 | 1000263 | 2.0% | |
| PC24A | 811064 | 800956 | 755206 | 5.5% | |
| PC24B | 891907 | 881539 | 832907 | 3.2% | |
| PC31A | 655794 | 647913 | 520477 | 12.2% | |
| PC41A | 2282146 | 2255293 | 1825864 | 1.5% | |
| PC41B | 1423437 | 1405744 | 1309049 | 1.2% | |
| PC43A | 295817 | 291848 | 185602 | 88.4% | Removed for high missing data |
| PC43B | 850190 | 839711 | 790031 | 2.7% | |
| PC43C | 942320 | 930580 | 870854 | 4.7% | |
| PK10 | 474110 | 468451 | 434443 | 10.2% | |
| PK11 | 514631 | 508454 | 479777 | 9.5% | |
| PK12 | 855470 | 844958 | 757425 | 4.0% | |
| PK13 | 389568 | 384551 | 351793 | 18.9% | |
| PK14 | 1461124 | 1442937 | 1336664 | 1.8% | |
| PK15 | 2091033 | 2064660 | 1910829 | 0.7% | |
| PK17 | 976629 | 963948 | 899350 | 3.4% | |
| PK18 | 245313 | 242304 | 226449 | 42.6% | |
| PK2 | 463925 | 458708 | 431899 | 10.7% | |
| PK3 | 657856 | 649643 | 610117 | 5.9% | |
| PK4 | 675886 | 667713 | 627699 | 6.9% | |
| PK5 | 896757 | 886463 | 819785 | 3.9% | |

| | | | | | |
|---|---|---|---|---|---|
| PK6_2 | 592971 | 585378 | 548619 | 8.6% | Concatenated with PK6 |
| PK6 | 458989 | 453001 | 426058 | 11.7% | |
| PK7 | 839749 | 829776 | 777617 | 4.2% | |
| PK8 | 766979 | 755601 | 704046 | 3.9% | |
| PK9 | 833848 | 821917 | 763002 | 4.6% | |
| PN10 | 344504 | 340424 | 317486 | 16.5% | |
| PN12 | 569181 | 561401 | 522964 | 6.7% | |
| PN13 | 711392 | 702180 | 609817 | 6.7% | |
| PN14 | 924666 | 914076 | 852939 | 3.1% | |
| PN15 | 464208 | 457889 | 423016 | 11.5% | |
| PN16 | 641685 | 634013 | 584908 | 6.0% | |
| PN18 | 1238751 | 1221463 | 1119397 | 2.4% | |
| PN1 | 877516 | 867613 | 806670 | 2.9% | |
| PN20 | 1069251 | 1054482 | 965214 | 2.5% | |
| PN21 | 763261 | 753921 | 695849 | 4.7% | |
| PN23 | 436129 | 430622 | 397612 | 12.1% | |
| PN24 | 198347 | 196198 | 183810 | 29.3% | |
| PN25 | 1596527 | 1576038 | 1453685 | 1.3% | |
| PN26 | 785646 | 776165 | 715041 | 3.8% | |
| PN28 | 803627 | 792829 | 739129 | 4.7% | |
| PN2 | 528804 | 522799 | 480437 | 8.3% | |
| PN5 | 1305647 | 1288594 | 1201312 | 1.7% | |
| PN6_2 | 291667 | 287899 | 262726 | 23.4% | Concatenated with PN6 |
| PN6 | 733024 | 723624 | 654781 | 5.0% | |

| | | | | |
|---|---|---|---|---|
| PN8 | 1242411 | 1228949 | 1122308 | 1.5% |
| PN9 | 1077325 | 1062341 | 989114 | 3.8% |
| Average | 693161 | 684282 | 630089 | |
| Standard Error | 28787 | 28422 | 26182 | |

**Table S3 |** Initial angsd filters used before filtering samples with missing data.

-uniqueOnly 1 -remove_bads 1  -minMapQ 20 -minQ 25 -dosnpstat 1 -doHWE 1 -sb_pval 1e-5 -hetbias_pval 1e-5 -skipTriallelic 1 -minInd 138 -snp_pval 1e-5 -minMaf 0.05 -setMinDepthInd 1

**Table S4.** Post-filtering angsd filters used on both species combined reference for outlier detection

-uniqueOnly 1 -remove_bads 1 -minMapQ 30 -minQ 35 -dosnpstat 1 -doHWE 1 -sb_pval 1e-3 -hetbias_pval 1e-3 -skipTriallelic 1 -maxHetFreq 0.5 -minInd 64 -snp_pval 1e-5 -minMaf 0.05"

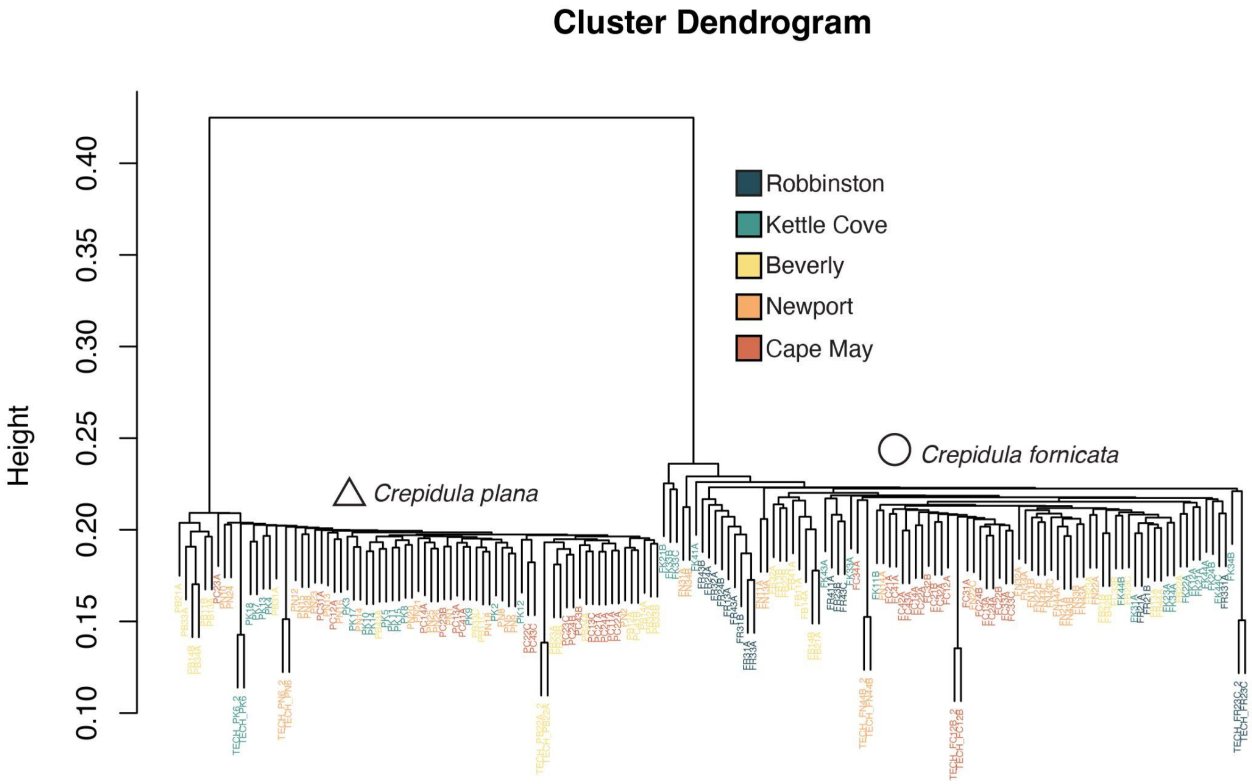

**Figure S1:** Hierarchical clustering of all samples. Technical replicates are highlighted with the tag TECH preceding sample names.

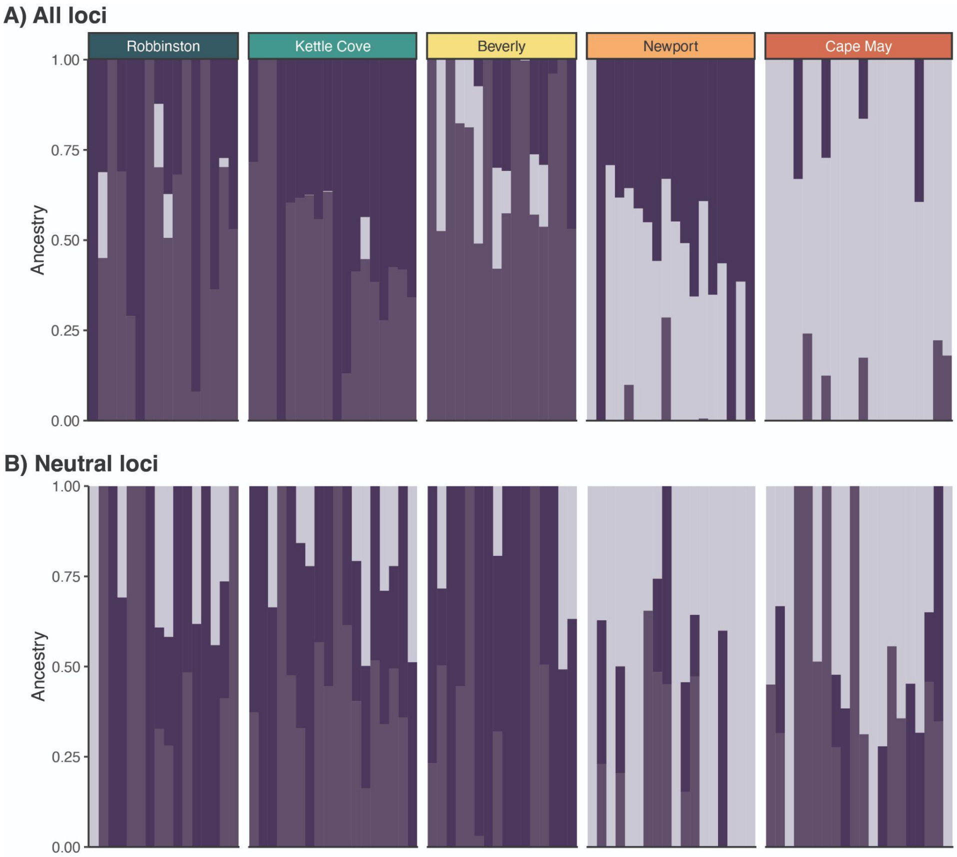

**Figure S2 |** ADMIXTURE results for *Crepidula fornicata* with optimal K = 3 determined using the Evanno method (Evanno et al., 2005) for A) all loci (N=4,379 SNPs) and B) neutral loci only (N=3,570 SNPs). Columns represent an individual sample and the bar color represents an individual's inferred membership into the three respective ancestral populations. Boxes above the data group individuals by collection sites.

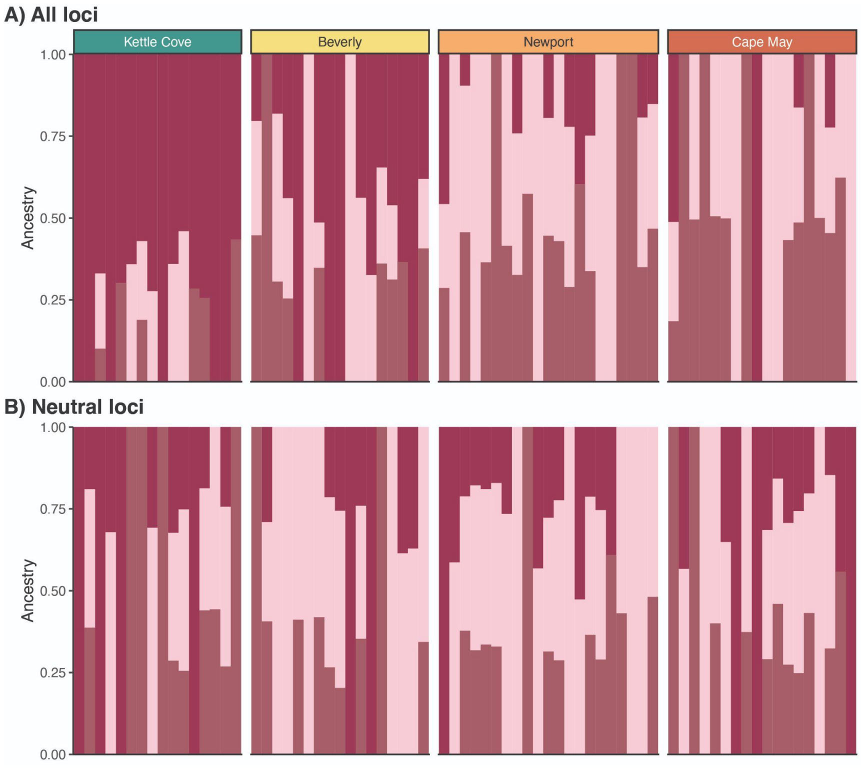

**Figure S3 |** ADMIXTURE results for *Crepidula plana* with optimal K = 3 determined using the Evanno method (Evanno et al., 2005) for A) all loci (N= 19,820 SNPs) and B) neutral loci only (N=17,322 SNPs). Columns represent an individual sample and the bar color represents an individual's inferred membership into the three respective ancestral populations. Boxes above the data group individuals by collection sites.

**Table S5 |** Pairwise genetic divergences (FST) between sites in A) *Crepidula fornicata* (purple) and B) *C. plana* (pink). Lower $F_{ST}$ values are shaded darker and higher $F_{ST}$ lighter to indicate degree of genetic divergence.

**A)**

| | Robbinston | Kettle Cove | Beverly | Newport |
|---|---|---|---|---|
| Kettle Cove | 0.010 | | | |
| Beverly | 0.009 | 0.010 | | |
| Newport | 0.010 | 0.011 | 0.009 | |
| Cape May | 0.017 | 0.016 | 0.012 | 0.007 |

**B)**

| | Kettle Cove | Beverly | Newport |
|---|---|---|---|
| Beverly | 0.007 | | |
| Newport | 0.013 | 0.007 | |
| Cape May | 0.014 | 0.009 | 0.009 |

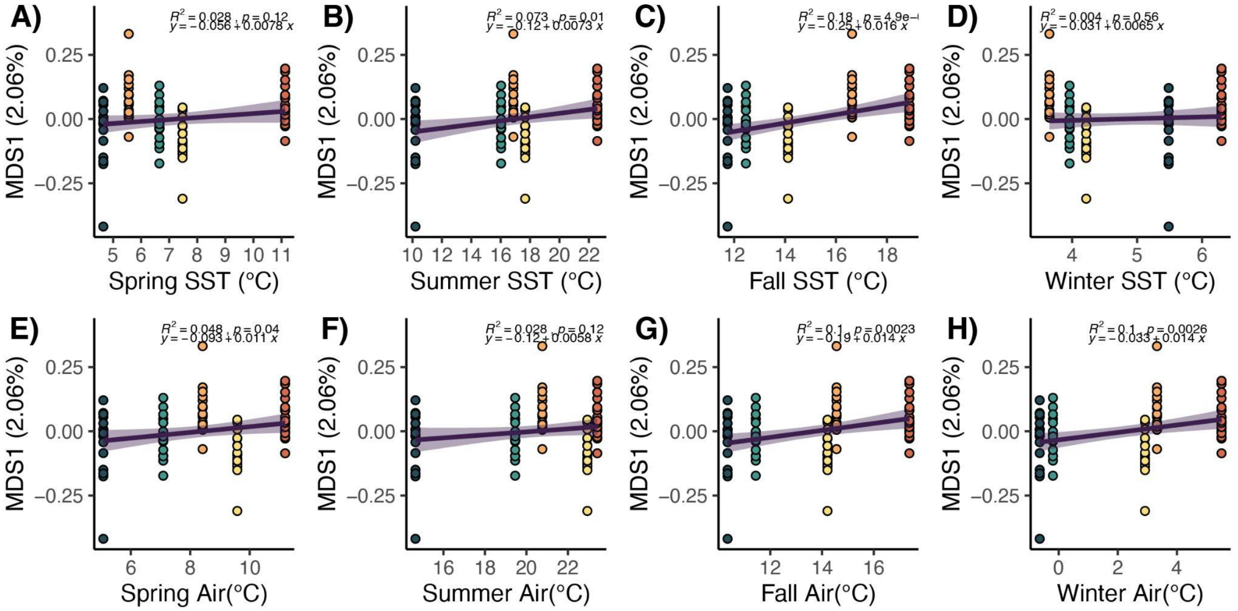

**Figure S4 |** Correlations between average seasonal water (A-D) and air temperatures (E-H) collected from NOAA weather buoys (See Table A3-1) with population genetic structure in neutral loci (N=3,570 SNPs) along the first MDS axis from in C. fornicata. Line of best fit is shown, which was calculated using a linear model, with 95% confidence intervals shaded. Note that direction of correlation is not informative due to the MDS1 axis being unitless.

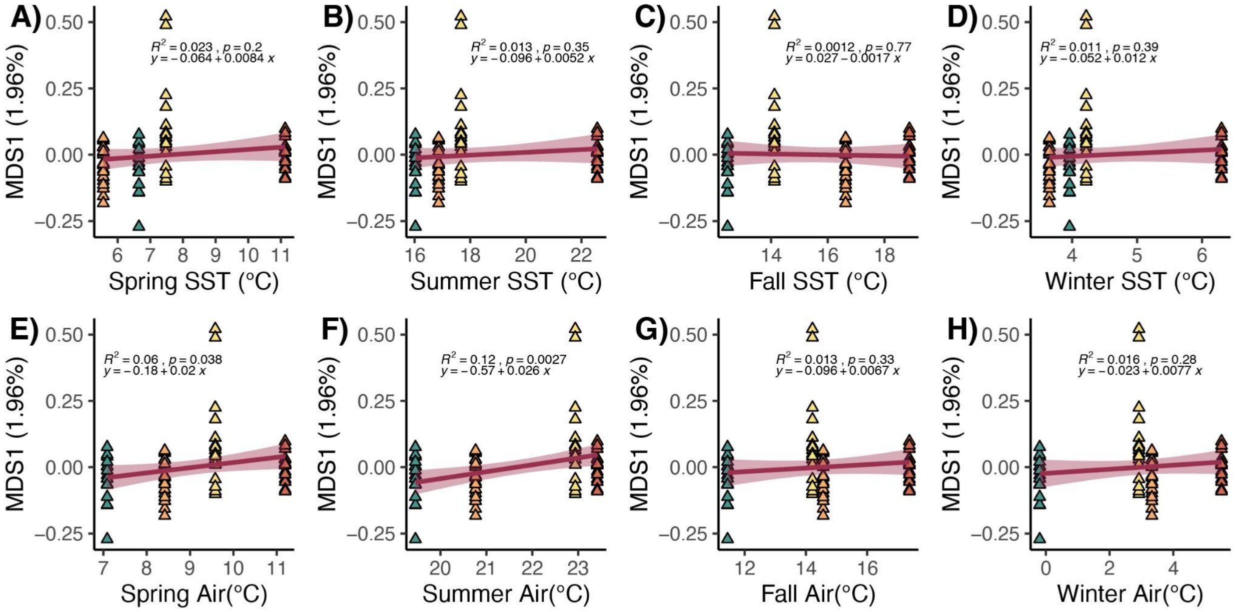

**Figure S5 |** Correlations between average seasonal water (A-D) and air temperatures (E-H) collected from NOAA weather buoys (See Table A3-1) with population genetic structure in neutral loci (N=17,322 SNPs) along the first MDS axis from Figure 4.4 1 in C. plana. Line of best fit is shown, which was calculated using a linear model, with 95% confidence intervals shaded. Note that direction of correlation is not informative due to the MDS1 axis being unitless.

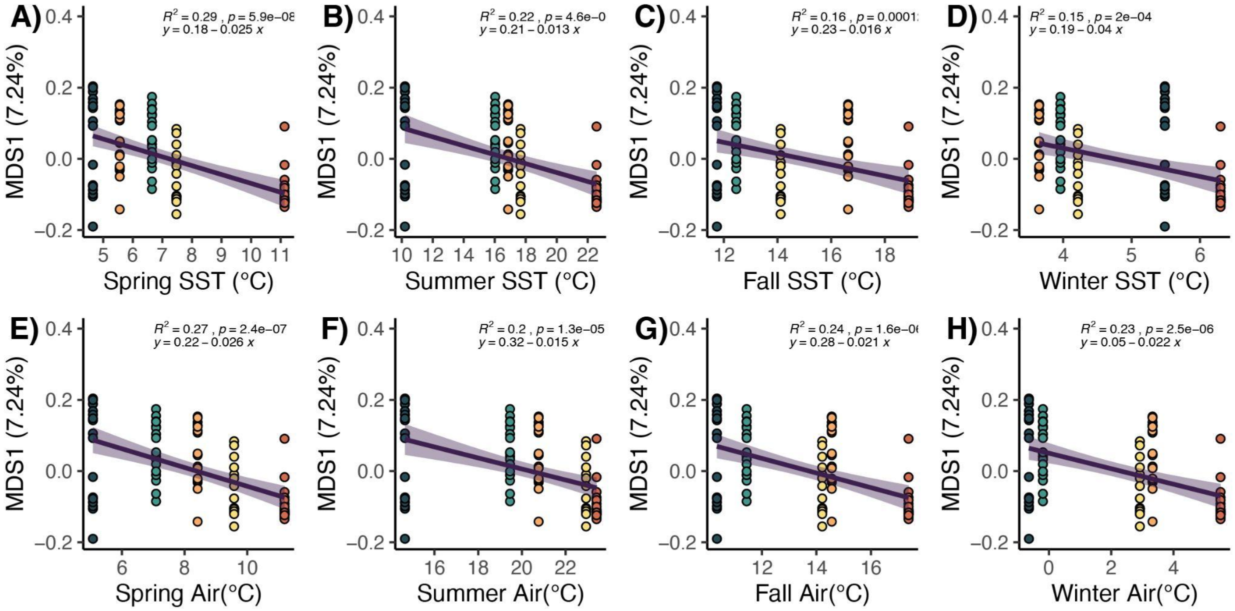

**Figure S6 |** Correlations between average seasonal water (A-D) and air temperatures (E-H) collected from NOAA weather buoys (See Table A3-1) with population genetic structure of outlier loci (N = 225 SNPs) along the first MDS axis from Figure 4.4 1 in *C. fornicata*. Linear models with lines of best fit with 95% confidence intervals shaded are shown. Note that direction of correlation is not informative due to the MDS1 axis being unitless.

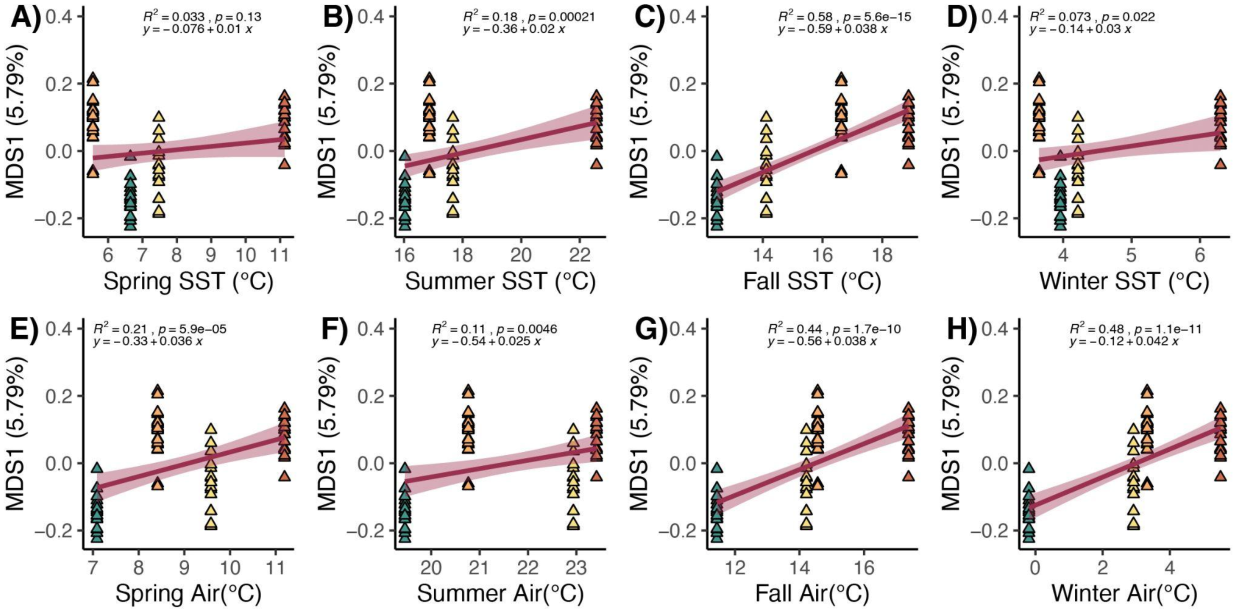

**Figure S7** | Correlations between average seasonal water (A-D) and air temperatures (E-H) collected from NOAA weather buoys (See Table A3-1) with population genetic structure of outlier loci (N = 317 SNPs) along the first MDS axis from Figure 4.4 1 in C. plana. Linear models with lines of best fit with 95% confidence intervals shaded are shown. Note that direction of correlation is not informative due to the MDS1 axis being unitless.
